# Increased expression of MCPIP1 in HIV-1 controllers is correlated with overexpression of p21

**DOI:** 10.1101/611871

**Authors:** Suwellen S. D. de Azevedo, Marcelo Ribeiro-Alves, Fernanda H. Côrtes, Edson Delatorre, Brenda Hoagland, Beatriz Grinsztejn, Valdilea G. Veloso, Mariza G. Morgado, Thiago Moreno L. Souza, Gonzalo Bello

**Affiliations:** Laboratório de AIDS & Imunologia Molecular. Instituto Oswaldo Cruz – IOC, FIOCRUZ. Rio de Janeiro, Brazil; Laboratório de Pesquisa Clínica em DST-AIDS. Instituto Nacional de Infectologia Evandro Chagas - INI, FIOCRUZ, Rio de Janeiro, Brazil; Laboratório de Genética Molecular de Microrganismos. Instituto Oswaldo Cruz – IOC, FIOCRUZ. Rio de Janeiro, Brazil; National Institute for Science and Technology on Innovation on Diseases of Neglected Populations (INCT/IDPN), Center for Technological Development in Health – CDTS, FIOCRUZ, Rio de Janeiro, Brazil; Laboratório de Imunofarmacologia. Instituto Oswaldo Cruz – IOC, FIOCRUZ. Rio de Janeiro, Brazil

**Author notes:** Correspondence: Suwellen S.D. de Azevedo /.

**Keywords:** HIV-1 controllers, restriction factors, MCPIP1, p21, immune activation

## Abstract

Some multifunctional cellular proteins, as the monocyte chemotactic protein-induced protein 1 (MCPIP1) and the cyclin-dependent kinase inhibitor p21, have also shown to be able to modulate the cellular susceptibility to the human immunodeficiency virus type 1 (HIV-1). Several studies described that p21 is expressed at high levels *ex vivo* in cells from individuals who naturally control HIV-1 replication (HIC). The expression level of MCPIP1 in HIC was never described before, but a recent study in a model of renal carcinoma cells showed that MCPIP1 overexpression was associated with an increase of both p21 transcripts and proteins levels. Here, we explored the potential associations between MCPIP1 and p21 expression, as well as with cellular activation in HIC, sustaining undetectable (elite controllers – EC) or low (viremic controllers – VC) viral loads. We found a selective upregulation of MCPIP1 and p21 mRNA levels in PBMC from HIC compared with both ART– suppressed and HIV–negative control groups (P ≤ 0.02) and a strong positive correlation (r ≥ 0.57; P ≤ 0.014) between expressions of both transcripts independently of the VL, treatment condition and HIV status. The mRNA levels of p21, but not of MCPIP1, were positively correlated with activated CD4^+^ T cells levels in HIC and EC (r ≥ 0.53; P ≤ 0.017). In relation to the monocyte activation, the mRNA levels of both p21 (r = 0.74; P = 0.005) and MCPIP1 (r = 0.58; P = 0.040) were positively correlated with plasmatic levels of sCD14 only in EC. Multivariate analysis confirmed the association between MCPIP1 and p21 mRNA levels, and between the latter with the frequency of activated CD4^+^ T cells. These data show for the first time the simultaneous overexpression and positive correlation of MCPIP1 and p21 transcripts in the setting of natural suppression of HIV-1 replication *in vivo*. The positive correlation between MCPIP1 and p21 transcripts supports a common regulatory pathway connecting these multifunctional host factors and a possible synergistic effect on HIV-1 replication control. Pharmacological manipulation of these cellular proteins may open novel therapeutic perspectives to prevent HIV-1 replication and disease progression.

## 1 Introduction

Among the individuals infected by the human immunodeficiency virus type 1 (HIV-1), a rare group called HIV controllers (HIC) suppress viral replication in absence of antiretroviral therapy, maintaining RNA viral loads (VL) below the limit of detection (LOD) (elite controllers, EC) or at low levels (> LOD and < 2,000 copies/ml; viremic controllers, VC). Natural control of HIV-1 replication is probably a multifactorial feature that involves different combinations of host and/or viral factors (1).

Some intrinsic host proteins, termed restriction factors (RF), are components of the innate immune response (2,3) that have the ability to cause a significant reduction in viral infectivity by interacting directly with the pathogen and are generally induced by interferon (IFN), hence being known as IFN-stimulated genes (ISGs) (4). Several RF has been shown to limit HIV replication *in vitro* at different stages of its life cycle (3), including some classical RF such the Apolipoprotein B mRNA-Editing enzyme, Catalytic polypeptide-like (APOBEC3G), the Bone Stromal Tumor protein 2 (BST2)/Tetherin, and the Sterile Alpha Motif domain and HD domain-containing protein 1 (SAMHD1) (2), and others more recently characterized like the Myxovirus resistance protein 2 (Mx2), the Interferon-inducible transmembrane family proteins (IFITM1-3 members) and Schlafen 11 (SLFN11) (3). The mRNA levels of some RF including SAMHD1, Theterin, IFITM1, Mx2 and SLFN11 have been described to be elevated in peripheral blood mononuclear cells (PBMC) or CD4^+^ T cells of HIC compared to antiretroviral (ART)-suppressed and/or HIV-uninfected individuals (5–9), although with contrasting findings across different HIC cohorts.

Others host multifunctional proteins, not recognized as classical RF, are also able to modulate the cellular susceptibility to HIV-1 infection. The cyclin-dependent kinase (CDK) inhibitor p21, encoded by the CDKN1A gene, modulates multiple relevant processes of the immune system, including proliferation of activated/memory T cells, macrophage activation and inflammation (10–17). This protein also indirectly limits the HIV-1 replication *in vitro* in various cellular systems by blocking the biosynthesis of dNTPs required for viral reverse transcription and by inhibiting the CDK9 activity required for HIV-1 mRNA transcription (18–23). Several studies described that p21 is expressed at high levels *ex vivo* in CD4^+^ T cells from HICs (21,24–26) and that p21 mRNA levels correlated with CD4^+^ T cell activation in EC, but not in other HIV-infected groups (5). These evidences suggest that the inducibility of p21 to immune activation is a singular characteristic of EC and may contribute to the natural control of HIV-1 replication *in vivo*.

The monocyte chemotactic protein–induced protein 1 (MCPIP1), encoded by ZC3H12A gene, is another newly discovered host multifunctional modulator of immune response with antiviral activity (27). MCPIP1 plays a critical role in the regulation of the inflammatory response and immune homeostasis and also blocks HIV-1 replication *in vitro* by promoting the viral mRNA degradation through its RNase activity, particularly in quiescent CD4^+^ T cells (27,28). In activated CD4^+^ T cells, MCPIP1 is rapidly degraded (28) after its cleavage by the mucosa-associated lymphoid-tissue lymphoma-translocation 1 (MALT1) protein (29,30). In activated macrophage cells, by contrast, MCPIP1 transcripts are induced by TLR ligands and pro-inflammatory cytokines (mainly, TNF-α, IL-1β and CCL2/MCP-1), and its expression stimulate a negative feedback loop that attenuates the inflammatory state by decreasing its fundamental mediators (27,31).

The expression level of MCPIP1 in HIC was never described before. Interestingly, a recent study in renal carcinoma cells (Caki-1 cells) revealed that MCPIP1 overexpression reduces the cellular growth by increasing the levels of p21 transcripts, along with other proteins involved in cell cycle progression/arrest, supporting a coordinate regulation of MCPIP1 and p21 transcripts in that cell-line (32). This evidence prompted us to ask whether the expression of MCPIP1 could be elevated and positively correlated with p21 in the setting of natural control of HIV-1 infection. To test this hypothesis, we quantified the *in vivo* expression of MCPIP1, p21 and several antiviral host RF mRNAs in PBMC from HIC, ART-suppressed and HIV-uninfected individuals. We further explored the potential relationship between MCPIP1/p21 expression and levels of systemic cellular activation in HIC.

## 2 Methods

### 2.1 Study Subjects

We analyzed a cohort of 21 HIC subjects followed-up at the Instituto Nacional de Infectologia Evandro Chagas (INI) in Rio de Janeiro, Brazil. All HIC maintained RNA VL of < 2,000 copies/ml without antiretroviral therapy for at least five years and were subdivided in two sub-groups: EC (*n* = 13) when most (≥ 70%) plasma VL determinations were below the limit of detection (LOD), and VC (*n* = 8) when most (≥ 70%) VL determinations were > LOD and < 2,000 copies/ml. The limit of detection of plasma VL determinations varied over the follow-up period in according to the Brazilian Ministry of Health guidelines, with methodologies being updated overtime to improve sensitivity: Nuclisens HIV-1 RNA QT assay (Organon Teknika, Durham, NC, limit of detection: 80 copies/mL) from 1999 to 2007; the Versant HIV-1 3.0 RNA assay (bDNA 3.0, Siemens, Tarrytown, NY, limit of detection: 50 copies/mL) from 2007 to 2013; and the Abbott RealTime HIV-1 assay (Abbott Laboratories, Wiesbaden, Germany, limit of detection: 40 copies/mL) from 2013 to until today. Virological and immunological characteristics of these subjects were described in detail in previous studies (33,34). Two groups of ART-suppressed subjects (ART, *n* = 8) and healthy HIV-1-uninfected subjects (NEG, *n* = 10) were used as controls.

### 2.2 mRNA gene-expression analysis

Total RNA was extracted from 1 × 10^7^ PBMC using RNeasy mini kit (Qiagen, Hilden, North Rhine-Westphalia, Germany) in which buffer RLT was supplemented with β-mercaptoethanol and displaced on-column DNase treatment using a Qiagen RNase-Free DNase Set (Qiagen, Hilden, North Rhine-Westphalia, Germany) according to manufacturer’s instruction. Total RNA yield and quality were determined using NanoDrop^®^ 8000 spectrophotometer and an Agilent^®^ 2100 Bioanalyzer. Only samples with an RNA integrity number (RIN) greater than 8.0 were used. Purified RNA (1 μg) was reverse-transcribed to cDNA using RT^2^ First Strand Kit (Qiagen, Hilden, North Rhine-Westphalia, Germany). The cDNA was mixed with RT^2^SYBR Green/ROX qPCR Master Mix (Qiagen, Hilden, North Rhine-Westphalia, Germany) and the mixture was added into customized RT^2^RNA PCR Array (Qiagen, Hilden, North Rhine-Westphalia, Germany) to measure the mRNA expression of 10 cellular target genes (APOBEC3G, SAMHD1, Tetherin, Mx1, Mx2, SLFN11, IFITM1, IFITM3, MCPIP1, and p21) besides three housekeeping genes (GAPDH, β-actin, and RNase-P), according to manufacturer’s instructions. Values of the crossing point at the maximum of the second derivative of the four-parameters fitted sigmoid curve second derivative, Cp, was determined for each sample. The efficiency of each amplification reaction was calculated as the ratio between the fluorescence of the cycle of quantification and fluorescence of the cycle immediately preceding that. Genes used in the normalization among samples were selected by the geNorm method (35). Data were expressed as fold-changes in mRNA abundance calculated as the normalized gene expression in any test sample divided by the mean normalized gene expression in the control HIV-negative group.

### 2.3 T cell and monocyte activation analyses

We used data of T cell and monocyte activation obtained in a previous study conducted by our group including these patients (34), in which plasma levels of soluble CD14 (sCD14) were determined by ELISA-sCD14 Quantikine assay (R&D Systems Minneapolis, MN) according to the manufacturer’s protocol and surface expression of combined HLA-DR and CD38 on CD4^+^ and CD8^+^ T cells was analyzed by flow cytometry.

### 2.4 Data analyses

The comparisons of mean log-fold changes in mRNA abundance were performed by either t-tests or one-way ANOVA nonparametric permutation tests (B = 1,000 permutations), followed by pair-wise comparisons with Holm-Bonferroni adjustment (36), for two or more groups respectively. Spearman coefficient was used for correlation analyses. A first-order log-Normal multiple regression analysis was fitted to model p21 gene expression as a function of MCPIP1 gene expression, CD4^+^ T cell activation (HLA-DR^+^CD38^+^), and HIC groups (EC and VC). The threshold for statistical significance was set to P < 0.05. Data were analyzed with R software (version 3.5.2) (37).

## 3 Results

Twenty-nine HIV-1 positive (21 HIC and 8 ART-suppressed) and 10 HIV-negative individuals were included in this cross-sectional study. Most HIV-positive (59%) and HIV-negative (60%) individuals were females and all individuals displayed CD4^+^ T cells counts above 500 cells/μl (Table 1). Although the EC subgroup shows a higher proportion of females (77%), the difference was not significant (Supplementary Table 1).

**Table 1.**
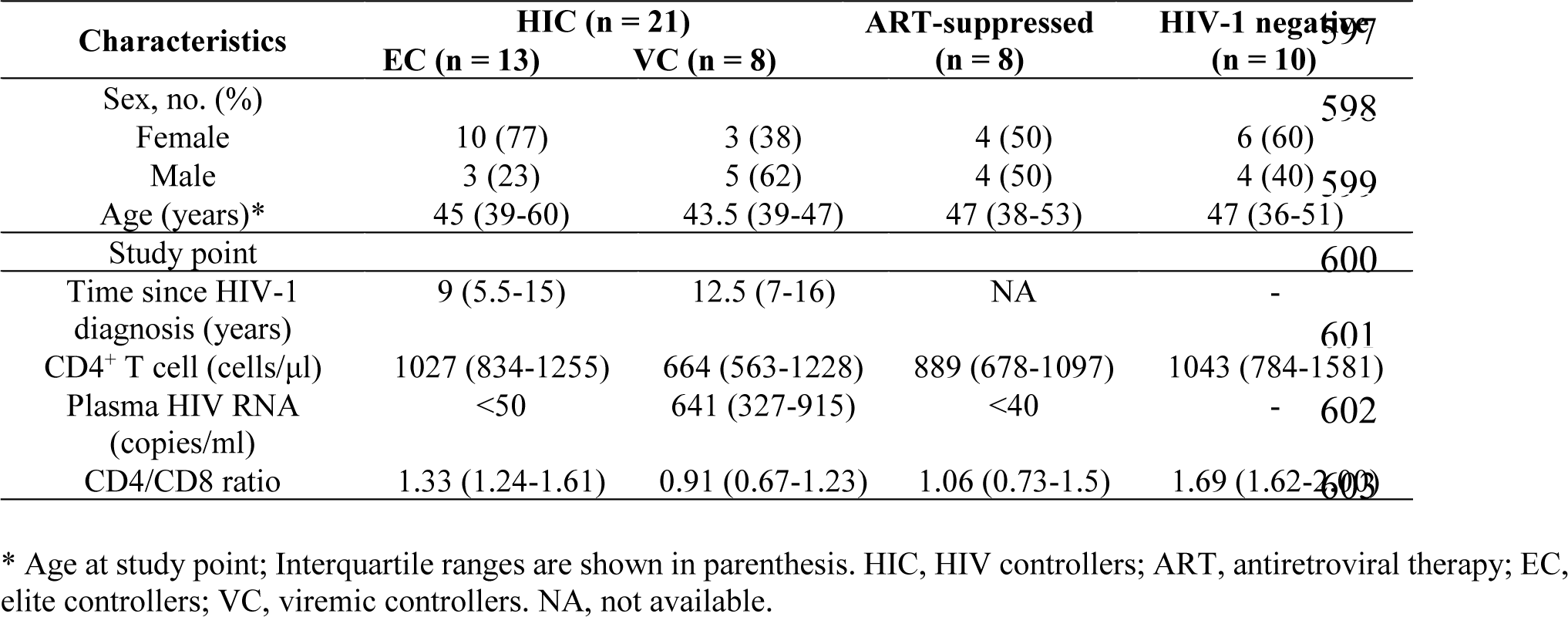
Main clinical and epidemiologic characteristics of individuals of this study.

| Characteristics | HIC (n = 21) |  | ART-suppressed<br>(n = 8) | HIV-1 negative<br>(n = 10) |
| --- | --- | --- | --- | --- |
|  | EC (n = 13) | VC (n = 8) |  |  |
| Sex, no. (%) |  |  |  |  |
| Female | 10 (77) | 3 (38) | 4 (50) | 6 (60) |
| Male | 3 (23) | 5 (62) | 4 (50) | 4 (40) |
| Age (years)* | 45 (39-60) | 43.5 (39-47) | 47 (38-53) | 47 (36-51) |
| Study point |  |  |  |  |
| Time since HIV-1<br>diagnosis (years) | 9 (5.5-15) | 12.5 (7-16) | NA | - |
| CD4<sup>+</sup> T cell (cells/μl) | 1027 (834-1255) | 664 (563-1228) | 889 (678-1097) | 1043 (784-1581) |
| Plasma HIV RNA<br>(copies/ml) | <50 | 641 (327-915) | <40 | - |
| CD4/CD8 ratio | 1.33 (1.24-1.61) | 0.91 (0.67-1.23) | 1.06 (0.73-1.5) | 1.69 (1.62-2.00) |

\* Age at study point; Interquartile ranges are shown in parenthesis. HIC, HIV controllers; ART, antiretroviral therapy; EC, elite controllers; VC, viremic controllers. NA, not available.

Analysis of the expression of multifunctional genes revealed a significant upregulation of both MCPIP1 and p21 transcripts in PBMC from HIC (Figure 1). The MCPIP1 mRNA was upregulated in PBMC from HIC compared to cells from both ART-suppressed (1.68-fold increase; P = 0.003) and HIV-negative (1.37-fold increase; P = 0.02) individuals (Figure 1A). A similar overexpression of the p21 mRNA was observed in PBMC from HIC compared to ART-suppressed (1.63-fold increase; P = 0.003) and HIV-negative (1.55-fold increase; P = 0.003) individuals (Figure 1B). In contrast, we found no significant differences in the mRNA levels of antiretroviral RF between the HIC and control groups, with the only exception of IFITM1 that was significantly elevated (1.15-fold increase; P = 0.03) in HIC in comparison to the HIV-negative group (Supplementary Figure S1).

**Figure 1.**
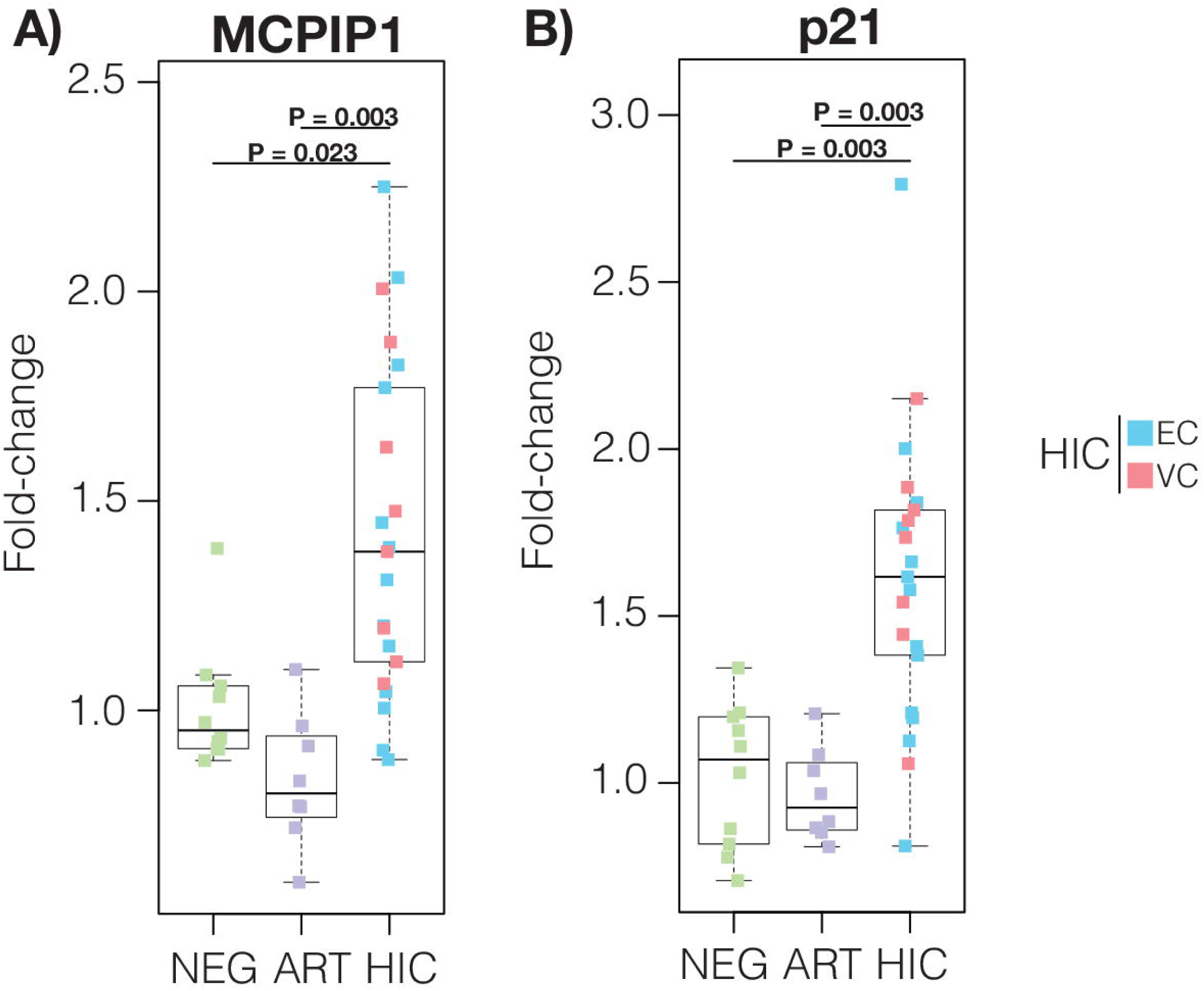
MCPIP1 and p21 mRNA levels are upregulated in PBMC from HIC. Boxplots represent the interquartile and sample median (central solid black line) of the relative changes (fold-change values relative to the mean of HIV-1-uninfected (NEG) subjects) of MCPIP1 (A) and p21 (B) expression comparing NEG and ART-suppressed subjects (ART) with HIV controllers (HIC). P-values < 0.05 were considered statistically significant.

We observed a significant positive correlation between the mRNA expression of MCPIP1 and p21 (r ≥ 0.57; *P* ≤ 0.014) in our cohort independently of the VL, treatment condition and HIV status (Figure 2). This positive correlation was maintained when individuals were subdivided by sex (Supplementary Figure S2). No significant correlations were observed between the mRNA expression of multifunctional genes MCPIP1/p21 and RF, with the only exception of a significant, negative correlation between MCPIP1/p21 and APOBEC3G in HIC (Supplementary Figure S3) and EC (Supplementary Figure S4).

**Figure 2.**
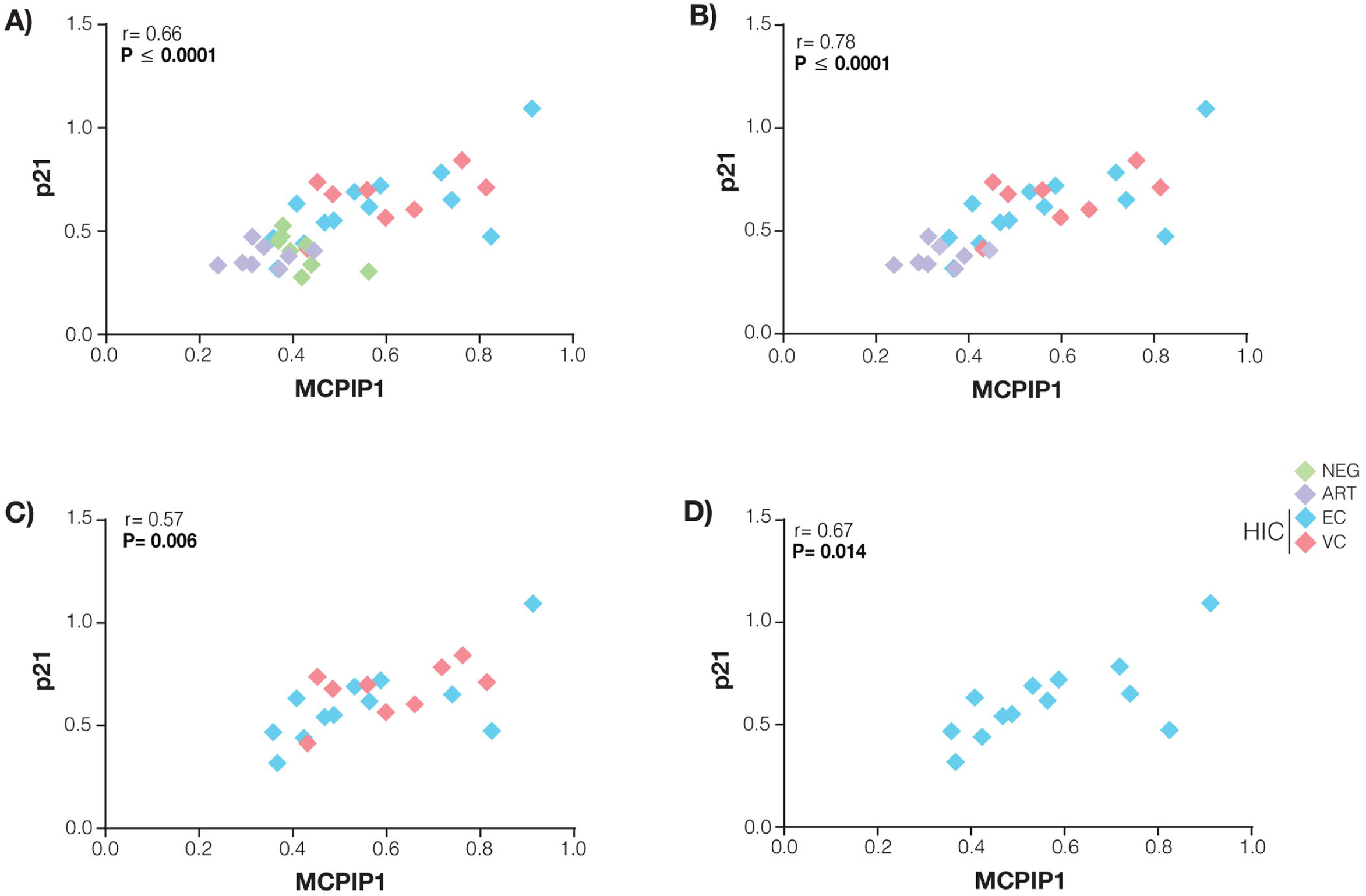
p21 and MCPIP1 mRNA levels in PBMC from HIC are positively correlated. The p21 and MCPIP1 normalized expression correlations were calculated considering all groups (A), HIV-infected (B), HIC (C), and EC (D). The points’ colors indicate the patient group, accordingly to the legend. Correlation coefficients (Spearman’s ρ) are shown in the upper right corner of each graph. P-values < 0.05 were considered statistically significant.

To explore the potential relationship of p21 or MCPIP1 expression with immune activation, we measured the frequency of phenotype HLA-DR^+^CD38^+^ on CD4^+^ and CD8^+^ T cells (T cell activation) and plasma levels of sCD14 (monocyte activation) in our cohort. Frequencies of activated CD4^+^ T cell populations in VC and ART-suppressed subjects were higher than in EC (P < 0.0001) and HIV-negative (P = 0.0002) individuals (Supplementary Figure S5A). The VC subgroup also had significantly higher frequencies of activated CD8^+^ T cell than EC (P = 0.0007) and control groups (P ≤ 0.0009) (Supplementary Figure S5B). The median concentration of sCD14 in plasma was not significantly different across the groups (Supplementary Figure S5C). No significant correlations between mRNA levels of MCPIP1 and CD4^+^ T cell (Figure 3A) or CD8^+^ T cell (data not shown) activation were observed for HIC or EC subsets. The mRNA levels of p21 were positively associated with activated CD4^+^ T cells levels in HIC (r = 0.53; P = 0.016) and EC (r = 0.68; P = 0.017) (Figure 3B); but not with activated CD8^+^ T cell levels (data not shown). Levels of sCD14 were positively correlated with both MCPIP1 (r = 0.58; P = 0.04) and p21 (r = 0.74; P = 0.005) mRNA levels only in the EC subset (Figure 3C and D). No significant correlations between mRNA levels of MCPIP1/p21 and CD4^+^/CD8^+^ T cell activation or sCD14 levels were observed when ART-suppressed and HIV-negative individuals were included (Supplementary Figures S6). Multivariate analysis showed that the upregulation of MCPIP1 was positively associated with the increase of p21 expression in HIC (1.44-fold increase; P = 0.0035) (Supplementary Figure S7A). The frequency of activated CD4^+^ T cells also was positively associated with the increase of p21 expression in both EC and VC (1.48-fold increase; P = 0.0116), although this increase of the p21 expression was down-regulated by the increase of activated CD4^+^ T cells in VC when compared to EC (1.30-fold decrease by an increase of 1% CD4^+^HLA-DR^+^CD38^+^ T cells; P = 0.0284) (Supplementary Figure S7B). Overall, the model was highly significant (P = 0.003) and could explain as much as 70% (R^2^ = 0.492) of p21 expression.

**Figure 3.**
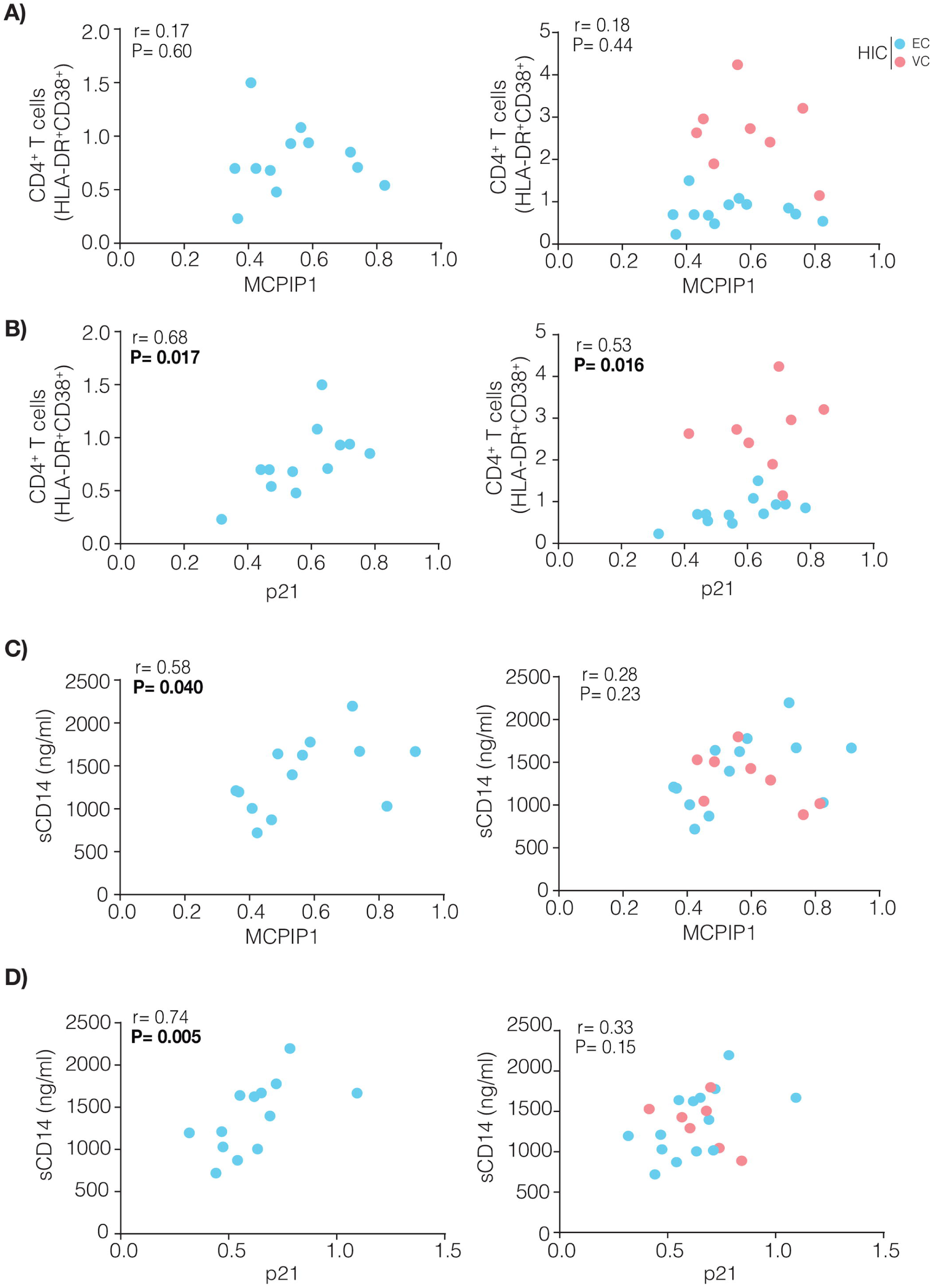
p21 transcripts are positively correlated with CD4^+^ T cell and monocyte activation while MCPIP1 transcripts are positively correlated only with monocyte activation in EC. The correlations were made evaluating the relationship between activated CD4^+^ T cells (A and B) or sCD14 levels (C and D) with the normalized expression of p21 and MCPIP1 for EC and HIC groups. The points’ colors present in each graph indicate the groups present according to the legend. Correlations coefficient (Spearman’s ρ) are shown in the upper left corner of each graph.

## 4 Discussion

In this study, we observed that MCPIP1 and p21 mRNA expression were significantly increased in PBMC of HIC compared to cells of HIV-negative and -positive/ART-suppressed individuals. While elevated expression of p21 in PBMC of HIC had already been previously described (5,21,24–26), this is the first study to show overexpression of MCPIP1 alongside with p21 in these individuals.

The mRNA levels of MCPIP1 and p21 were positively correlated in HIC as well as in HIV–positive and –negative individuals. This supports a coordinated expression of these cellular genes in different settings, consistent with what has been shown for a renal carcinoma cell line (32). According to this study, MCPIP1 expression triggers the activation of p21 by two mechanisms: 1) down-modulation of damage-specific DNA binding protein 1 (DDB1) which regulates degradation of p21; and 2) upregulation of the mRNA levels of chromatin licensing and DNA replication factor 1 (CDT1) which activates p21 (32). In addition, following HIV-1 infection, the cellular let-7c miRNA is upregulated and it downregulate p21, resulting in higher copy number of viral genome transcripts in infected cells (38). MCPIP1 acts as a broad suppressor of the biogenesis pathway of both cellular (39) and viral miRNA (40). The involvement of the MCPIP1 in the degradation of another precursor of let-7 family (pre-let-7g) was already described (41), reinforcing the hypothesis that MCPIP1 might enhance the antiviral responses triggered by HIV-1 entry and infection by downregulating the miRNAs that target p21.

Increased expression of some host RF, which are also ISGs (4), has been previously observed in CD4^+^ T cells (i.e., SAMHD1, SLFN11 and IFITM1) (5,7,8) and PBMC (i.e., Mx1, Mx2, Tetherin and SLFN11) from HIC (6,9). With the only exception of IFITM1, no other RF analyzed here were upregulated in PBMC from our HIC cohort. In the chronic phase of HIV-1 infection in viremic untreated patients, most ISGs are upregulated in CD4^+^ T cells (42–44) and their expression is positively correlated with the percentage of activated T cells and negatively correlated with CD4^+^ T cell counts (42–46). This suggests that residual or low-level viremia observed in our HIC might not be enough to induce a generalized upregulation of ISGs during chronic infection (44). In addition, MCPIP1 (47,48) and p21 (16) negatively regulate the NF-κB cascade and their overexpression may also contribute to limit the chronic overexpression of ISGs in HIC. While most RF are mainly induced by IFN type I, IFITM1 can also be induced by IFN type II (49), indicating that another pathway may have stimulated its expression in our HIC cohort.

Although we have failed to detect an overall up-regulation of host RF in our HIC cohort, it is interesting to note that a few individuals displayed mRNA levels of SAMHD1 and/or SLFN11 well above the normal range (Supplementary Figure S1). These observations suggest that there might not be a unique host RF expression signature common to all HIC, but that different combinations of host RF could be associated with natural control of HIV-1 replication in distinct individuals. Thus, the particular set of increased host RF may vary across different HIC cohorts and this might explain the apparently contrasting findings across studies (5–9,50). Additionally, even though we were able to identify statistically significant differences in expression levels of MCPIP1 and p21 in PBMC between HIC and control groups, these findings warrant validation using larger cohorts.

Our results confirm previous observations that levels of p21 mRNA are positively correlated with CD4^+^ T cell activation in EC and HIC groups (5) and further support a positive correlation with sCD14, a marker of monocyte activation, in EC. These correlations are fully consistent with the critical role of p21 as a negative regulator of the proliferation of activated/memory T cells (10,13,14) and of macrophage-mediated inflammatory responses (15–17). Although MCPIP1 expression is also essential for suppressing peripheral T cell (51) and macrophage (52,53) activation, we only found a positive correlation of MCPIP1 mRNA with sCD14 in EC. While induction of MCPIP1 mRNA *in vitro* in response to TLR as well as IL-1β stimulation in macrophages is rapid and long-lasting (≥ 24h) (52– 54), the corresponding induction upon T cell receptor stimulation in CD4^+^ T cell is more ephemeral (< 12 hours) (55), which could have hindered the observation of a direct correlation between these two parameters. Notably, increased expression of MCPIP1/p21 associated with T cell and/or monocyte activation seems to be a unique characteristic of HIC/EC, because similar correlations were not observed in our study for other HIV-infected or HIV-negative subjects and previous studies have shown that viremic progressors display reduced levels of p21 even though exhibit high levels of cellular activation and inflammation (21). These results suggest that MCPIP1/p21 overexpression may be a distinctive homoeostatic innate response of HIC to limit the deleterious effects of aberrant chronic immune activation and inflammation driven by HIV-1 infection.

Transcript levels of RF here analyzed were not significantly correlated with T cell activation or sCD14, with the only exception of a negative correlation between APOBEC3G mRNA and sCD14 levels in EC (r = - 0.73. P = 0.006; data not shown). Surprisingly, transcripts levels of APOBEC3G were also negatively correlated with MCPIP1 and p21 mRNA levels in both HIC and EC. One possible explanation for these negative correlations lies in the interaction of APOBEC3G, MCPIP1, and p21 with the product of an important monocyte differentiation gene, the Kruppel-like factor 4 (KLF4). The expression of KLF4 in human macrophages is induced after IFN-γ, LPS, or TNF-α stimulus (56), mediating the proinflammatory signaling and the direct transcriptional regulation of CD14 *in vitro* (57). Interestingly, KLF4 is also able to induce expression of both MCPIP1 (58) and p21 (59,60), whereas APOBEC3G binds to the 3’-UTR of KLF4 mRNA and results in the reduction of its expression (61). Thus, lower levels of APOBEC3G mRNA may be associated with an upregulation of KLF4 that in turn induce higher levels of sCD14 and MCPIP1/p21 mRNA.

Selective upregulation of MCPIP1 and p21 in CD4^+^ T, macrophages and/or dendritic cells may directly limit HIV-1 replication by 1) reducing the reverse transcription and chromosomal integration of HIV-1 in quiescent cells and thus limiting the size of the latent proviral reservoir (18–20,62–64); 2) restricting HIV-1 LTR transcription (47,48,65,66); and, 3) degrading viral mRNA and miRNA (28,39,40,67). Upregulation of p21 and MCPIP1 may also indirectly limit HIV-1 replication and further prevent CD4^+^ T cells loss by reducing chronic IFN-I signaling, generalized inflammation and over-activation of the immune system (10,14–17,52,53,68–70), without affecting the activation of antiviral cellular responses. Although the enhanced antiviral and anti-inflammatory state may not be enough to fully restrict HIV-1 replication (71), it could act in concert with other innate and adaptive immune mechanisms to control HIV replication in HIC.

The enhanced expression of a few select host genes, including p21, was strongly associated with reduced CD4^+^ T cell-associated HIV RNA during ART, indicating that the p21 may contribute to the control of viral expression and ongoing replication during ART (72). Another study demonstrates that atorvastatin, a lipid-lowering medication, exert a broad spectrum of anti-inflammatory functions and further reduced HIV infection in both rested and activated CD4^+^ T cells *in vitro* via p21 upregulation (22). Interestingly, atorvastatin was found to up-regulates p21 through a p53 independent pathway, which is consistent with a potential role of MCPIP1 in that antiviral mechanism. These observations suggest that pharmacological manipulation of p21 and MCPIP1 may open novel therapeutic perspectives to prevent HIV-1 replication and to attenuate HIV-associated inflammation and immune activation during ART.

An important limitation of our study is the impossibility of assigning which cell(s) population(s) has increased expression of p21 and MCPIP1 in HIC. The expression profile of many RF and ISGs may be different between CD4^+^ T cells and monocytes (8), suggesting that the individualization of these cell types might better decipher the mechanisms of host factors regulation in the setting of natural control of HIV-1 infection. Another potential limitation is that only mRNA levels were analyzed. Previous studies showed that p21 mRNA levels mirror p21 protein levels in CD4^+^ T cells from HIC (21) and that MCPIP1 mRNA levels reflect MCPIP1 protein levels in HCV-infected hepatoma cells (73). Although this evidence indicates a close match between transcripts and protein expression levels, measuring the levels/activity of p21 and MCPIP1 proteins in cells from HIC should also help to elucidate the relevance of these RF for HIV control.

In summary, our data confirm the high levels of p21 mRNA expression and shows for the first-time the concurrent overexpression of MCPIP1 mRNA in HIC. Moreover, we found a positive correlation between p21 and MCPIP1 transcripts in HIC, indicating a possible synergistic effect of both innate host RF on natural suppression of HIV-1 replication *in vivo*. Further studies are needed to better understand the role of p21 and MCPIP1 in the natural control of HIV-1 replication and disease progression in HIC. These findings may also have important implications for the development of new immune-based therapeutic strategies for a functional cure of HIV-1 infection.

## 6 Ethics Statement

This study was carried out in accordance with the recommendations of the ethical committee of Instituto Nacional de Infectologia Evandro Chagas (INI-Fiocruz) that approved the study protocol (CAAE 1717.0.000.009-07). All subjects gave written informed consent in accordance with the Declaration of Helsinki.

## 8 Author Contributions

GB and TMLS conceived and designed the study and supervised the experiments. SSDA conducted experiments and analyzed the data together with MR-A and GB. FH performed the CD4^+^ T cell and monocyte activation assays. ED collaborated with mRNA gene-expression analysis. BH, BG, and VGV conducted patient recruitment and follow-up. FH, ED and MGM provided intellectual input for results interpretations. SSDA, GB and MR-A wrote the first draft and all authors assisted with the writing and approved the final manuscript.

## 9 Acknowledgments

The authors thank the patients, who participated in the study, as well as all the technical staff involved in the clinical follow-up of these patients. We also thank Ms Marilia Alves Figueira de Melo for the excellent technical support in RNA quantification and integrity analyses and the Plataforma de PCR em Tempo Real – RJ (RPT09A) – FIOCRUZ and Plataforma de Sequenciamento de Ácidos Nucléicos de Nova Geração – RJ (RPT01J) – FIOCRUZ.

## References

1. Walker BD, Yu XG. Unravelling the mechanisms of durable control of HIV-1. Nat Rev Immunol (2013) 13:487–498. doi:10.1038/nri3478

2. Harris RS, Hultquist JF, Evans DT. The Restriction Factors of Human Immunodeficiency Virus. J Biol Chem (2012) 287:40875–40883. doi:10.1074/jbc.R112.416925

3. Colomer-Lluch M, Ruiz A, Moris A, Prado JG. Restriction Factors: From Intrinsic Viral Restriction to Shaping Cellular Immunity Against HIV-1. Front Immunol (2018) 9:2876. doi:10.3389/fimmu.2018.02876

4. Doyle T, Goujon C, Malim MH. HIV-1 and interferons: who’s interfering with whom? Nat Rev Microbiol (2015) 13:403–413. doi:10.1038/nrmicro3449

5. Abdel-Mohsen M, Raposo RAS, Deng X, Li M, Liegler T, Sinclair E, Salama MS, Ghanem HEA, Hoh R, Wong JK, et al. Expression profile of host restriction factors in HIV-1 elite controllers. Retrovirology (2013) 10:1–13. doi:10.1186/1742-4690-10-106

6. Krishnan S, Wilson EMP, Sheikh V, Rupert A, Mendoza D, Yang J, Lempicki R, Migueles SA, Sereti I. Evidence for innate immune system activation in HIV type 1-infected elite controllers. J Infect Dis (2014) 209:931–939. doi:10.1093/infdis/jit581

7. Riveira-Muñoz E, Ruiz A, Pauls E, Permanyer M, Badia R, Mothe B, Crespo M, Clotet B, Brander C, Ballana E, et al. Increased expression of SAMHD1 in a subset of HIV-1 elite controllers. J Antimicrob Chemother (2014) 69:3057–3060. doi:10.1093/jac/dku276

8. Canoui E, Noël N, Lécuroux C, Boufassa F, Sáez-Cirión A, Bourgeois C, Lambotte O, ANRS CO21 CODEX Study Group. Strong ifitm1 Expression in CD4 T Cells in HIV Controllers Is Correlated With Immune Activation. J Acquir Immune Defic Syndr (2017) 74:e56–e59. doi:10.1097/QAI.0000000000001166

9. Van Hecke C, Trypsteen W, Malatinkova E, De Spiegelaere W, Vervisch K, Rutsaert S, Kinloch-de Loes S, Sips M, Vandekerckhove L. Early treated HIV-1 positive individuals demonstrate similar restriction factor expression profile as long-term non-progressors. EBioMedicine (2019) 1–12. doi:10.1016/j.ebiom.2019.02.006

10. Khanna AK, Plummer M, Nilakantan V, Pieper GM. Recombinant p21 Protein Inhibits Lymphocyte Proliferation and Transcription Factors. J Immunol (2005) 174:7610–7617. doi:10.4049/jimmunol.174.12.7610

11. Khanna AK. Reciprocal role of cyclins and cyclin kinase inhibitor p21WAF1/CIP1 on lymphocyte proliferation, allo-immune activation and inflammation. BMC Immunol (2005) 6:22. doi:10.1186/1471-2172-6-22

12. Balomenos D, Martín-Caballero J, García MI, Prieto I, Flores JM, Serrano M, Martínez-A C. The cell cycle inhibitor p21 controls T-cell proliferation and sex-linked lupus development. Nat Med (2000) 6:171–6. doi:10.1038/72272

13. Arias CF, Ballesteros-Tato A, Garcia MI, Martin-Caballero J, Flores JM, Martinez-A C, Balomenos D. p21CIP1/WAF1 Controls Proliferation of Activated/Memory T Cells and Affects Homeostasis and Memory T Cell Responses. J Immunol (2007) 178:2296–2306. doi:10.4049/jimmunol.178.4.2296

14. Santiago-Raber M-L, Lawson BR, Dummer W, Barnhouse M, Koundouris S, Wilson CB, Kono DH, Theofilopoulos AN. Role of Cyclin Kinase Inhibitor p21 in Systemic Autoimmunity. J Immunol (2001) 167:4067–4074. doi:10.4049/jimmunol.167.7.4067

15. Lloberas J, Celada A. p21 waf1/CIP1, a CDK inhibitor and a negative feedback system that controls macrophage activation. Eur J Immunol (2009) 39:691–694. doi:10.1002/eji.200939262

16. Trakala M, Arias CF, García MI, Moreno-Ortiz MC, Tsilingiri K, Fernández PJ, Mellado M, Díaz-Meco MT, Moscat J, Serrano M, et al. Regulation of macrophage activation and septic shock susceptibility via p21(WAF1/CIP1). Eur J Immunol (2009) 39:810–819. doi:10.1002/eji.200838676

17. Scatizzi JC, Mavers M, Hutcheson J, Young B, Shi B, Pope RM, Ruderman EM, Samways DSK, Corbett JA, Egan TM, et al. The CDK domain of p21 is a suppressor of IL-1β-mediated inflammation in activated macrophages. Eur J Immunol (2009) 39:820–825. doi:10.1002/eji.200838683

18. Zhang J, Scadden DT, Crumpacker CS. Primitive hematopoietic cells resist HIV-1 infection via p21. J Clin Invest (2007) 117:473–81. doi:10.1172/JCI28971

19. Bergamaschi A, David A, Le Rouzic E, Nisole S, Barre-Sinoussi F, Pancino G. The CDK Inhibitor p21Cip1/WAF1 Is Induced by Fc R Activation and Restricts the Replication of Human Immunodeficiency Virus Type 1 and Related Primate Lentiviruses in Human Macrophages. J Virol (2009) 83:12253–12265. doi:10.1128/JVI.01395-09

20. Valle-Casuso JC, Allouch A, David A, Lenzi GM, Studdard L, Barré-Sinoussi F, Müller-Trutwin M, Kim B, Pancino G, Sáez-Cirión A. p21 Restricts HIV-1 in Monocyte-Derived Dendritic Cells through the Reduction of Deoxynucleoside Triphosphate Biosynthesis and Regulation of SAMHD1 Antiviral Activity. J Virol (2017) 91:1–18. doi:10.1128/JVI.01324-17

21. Chen H, Li C, Huang J, Cung T, Seiss K, Beamon J, Carrington MF, Porter LC, Burke PS, Yang Y, et al. CD4+ T cells from elite controllers resist HIV-1 infection by selective upregulation of p21. J Clin Invest (2011) 121:1549–1560. doi:10.1172/JCI44539

22. Elahi S, Weiss RH, Merani S. Atorvastatin restricts HIV replication in CD4+ T cells by upregulation of p21. Aids (2016) 30:171–183. doi:10.1097/QAD.0000000000000917

23. Elahi S, Niki T, Hirashima M, Horton H. Galectin-9 binding to Tim-3 renders activated human CD4+ T cells less susceptible to HIV-1 infection. Blood (2012) 119:4192–4204. doi:10.1182/blood-2011-11-389585

24. Saez-Cirion A, Hamimi C, Bergamaschi A, David A, Versmisse P, Melard A, Boufassa F, Barre-Sinoussi F, Lambotte O, Rouzioux C, et al. Restriction of HIV-1 replication in macrophages and CD4+ T cells from HIV controllers. Blood (2011) 118:955–964. doi:10.1182/blood-2010-12-327106

25. Moosa Y, Tanko RF, Ramsuran V, Singh R, Madzivhandila M, Yende-Zuma N, Abrahams MR, Selhorst P, Gounder K, Moore PL, et al. Case report: Mechanisms of HIV elite control in two African women. BMC Infect Dis (2018) 18:1–7. doi:10.1186/s12879-018-2961-8

26. Madlala P, Van de Velde P, Van Remoortel B, Vets S, Van Wijngaerden E, Van Laethem K, Gijsbers R, Schrijvers R, Debyser Z. Analysis of ex vivo HIV-1 infection in a controller-discordant couple. J virus Erad (2018) 4:170–173.

27. Fu M, Blackshear PJ. RNA-binding proteins in immune regulation: a focus on CCCH zinc finger proteins. Nat Rev Immunol (2017) 17:130–143. doi:10.1038/nri.2016.129

28. Liu S, Qiu C, Miao R, Zhou J, Lee A, Liu B, Lester SN, Fu W, Zhu L, Zhang L, et al. MCPIP1 restricts HIV infection and is rapidly degraded in activated CD4+ T cells. Proc Natl Acad Sci U S A (2013) 110:19083–8. doi:10.1073/pnas.1316208110

29. Uehata T, Iwasaki H, Vandenbon A, Matsushita K, Hernandez-cuellar E, Kuniyoshi K, Satoh T, Mino T, Suzuki Y, Standley DM, et al. Malt1-Induced Cleavage of Regnase-1 in CD4 + Helper T Cells Regulates Immune Activation. Cell (2013) 153:1036–1049. doi:10.1016/j.cell.2013.04.034

30. Jeltsch KM, Hu D, Brenner S, Zöller J, Heinz GA, Nagel D, Vogel KU, Rehage N, Warth SC, Edelmann SL, et al. Cleavage of roquin and regnase-1 by the paracaspase MALT1 releases their cooperatively repressed targets to promote TH17 differentiation. Nat Immunol (2014) 15:1079–1089. doi:10.1038/ni.3008

31. Jura J, Skalniak L, Koj A. Monocyte chemotactic protein-1-induced protein-1 (MCPIP1) is a novel multifunctional modulator of inflammatory reactions. Biochim Biophys Acta (2012) 1823:1905–13. doi:10.1016/j.bbamcr.2012.06.029

32. Lichawska-Cieslar A, Pietrzycka R, Ligeza J, Kulecka M, Paziewska A, Kalita A, Dolicka DD, Wilamowski M, Miekus K, Ostrowski J, et al. RNA sequencing reveals widespread transcriptome changes in a renal carcinoma cell line. Oncotarget (2018) 9:8597–8613. doi:10.18632/oncotarget.24269

33. Azevedo SSD, Caetano DG, Côrtes FH, Teixeira SLM, Santos Silva K, Hoagland B, Grinsztejn B, Veloso VG, Morgado MG, Bello G. Highly divergent patterns of genetic diversity and evolution in proviral quasispecies from HIV controllers. Retrovirology (2017) 14:1–13. doi:10.1186/s12977-017-0354-5

34. Côrtes FH, de Paula HHS, Bello G, Ribeiro-Alves M, de Azevedo SSD, Caetano DG, Teixeira SLM, Hoagland B, Grinsztejn B, Veloso VG, et al. Plasmatic levels of IL-18, IP-10, and *A.*activated CD8+T cells are potential biomarkers to identify HIV-1 elite controllers with a true functional cure profile. Front Immunol (2018) 9:1–11. doi:10.3389/fimmu.2018.01576

35. Vandesompele J, De Preter K, Pattyn F, Poppe B, Van Roy N, De Paepe A, Speleman F. Accurate normalization of real-time quantitative RT-PCR data by geometric averaging of multiple internal control genes. Genome Biol (2002) 3:RESEARCH0034. doi:10.1186/gb-2002-3-7-research0034

36. Basso D, Pesarin F, Salmaso L, Solari A. “Nonparametric One-Way ANOVA”, in Permutation Tests for Stochastic Ordering and ANOVA (New York, NY: Springer), 105–132. doi:10.1007/978-0-387-85956-9_5

37. R Core Team. R: A language and environment for statistical computing., org. R Foundation for Statistical Computing Vienna, Austria. (2018).

38. Farberov L, Herzig E, Modai S, Isakov O, Hizi A, Shomron N. MicroRNA-mediated regulation of p21 and TASK1 cellular restriction factors enhances HIV-1 infection. J Cell Sci (2015) 128:1607–1616. doi:10.1242/jcs.167817

39. Suzuki HI, Arase M, Matsuyama H, Choi YL, Ueno T, Mano H, Sugimoto K, Miyazono K. MCPIP1 ribonuclease antagonizes dicer and terminates microRNA biogenesis through precursor microRNA degradation. Mol Cell (2011) 44:424–436. doi:10.1016/j.molcel.2011.09.012

40. Happel C, Ramalingam D, Ziegelbauer JM. Virus-Mediated Alterations in miRNA Factors and Degradation of Viral miRNAs by MCPIP1. PLoS Biol (2016) 14:e2000998. doi:10.1371/journal.pbio.2000998

41. Suzuki HI, Katsura A, Miyazono K. A role of uridylation pathway for blockade of let-7 microRNA biogenesis by Lin28B. Cancer Sci (2015) 106:1174–1181. doi:10.1111/cas.12721

42. Sedaghat AR, German J, Teslovich TM, Cofrancesco J, Jie CC, Talbot CC, Siliciano RF. Chronic CD4+ T-Cell Activation and Depletion in Human Immunodeficiency Virus Type 1 Infection: Type I Interferon-Mediated Disruption of T-Cell Dynamics. J Virol (2008) 82:1870–1883. doi:10.1128/JVI.02228-07

43. Hyrcza MD, Kovacs C, Loutfy M, Halpenny R, Heisler L, Yang S, Wilkins O, Ostrowski M, Der SD. Distinct Transcriptional Profiles in Ex Vivo CD4+ and CD8+ T Cells Are Established Early in Human Immunodeficiency Virus Type 1 Infection and Are Characterized by a Chronic Interferon Response as Well as Extensive Transcriptional Changes in CD8+ T Cells. J Virol (2007) 81:3477–3486. doi:10.1128/JVI.01552-06

44. Rotger M, Dang KK, Fellay J, Heinzen EL, Feng S, Descombes P, Shianna K V., Ge D, Günthard HF, Goldstein DB, et al. Genome-Wide mRNA Expression Correlates of Viral Control in CD4+ T-Cells from HIV-1-Infected Individuals. PLoS Pathog (2010) 6:e1000781. doi:10.1371/journal.ppat.1000781

45. Hardy GAD, Sieg S, Rodriguez B, Anthony D, Asaad R, Jiang W, Mudd J, Schacker T, Funderburg NT, Pilch-Cooper HA, et al. Interferon-α Is the Primary Plasma Type-I IFN in HIV-1 Infection and Correlates with Immune Activation and Disease Markers. PLoS One A.(2013) 8:e56527. doi:10.1371/journal.pone.0056527

46. Fernandez S, Tanaskovic S, Helbig K, Rajasuriar R, Kramski M, Murray JM, Beard M, Purcell D, Lewin SR, Price P, et al. CD4+ T-Cell Deficiency in HIV Patients Responding to Antiretroviral Therapy Is Associated With Increased Expression of Interferon-Stimulated Genes in CD4+ T Cells. J Infect Dis (2011) 204:1927–1935. doi:10.1093/infdis/jir659

47. Skalniak L, Mizgalska D, Zarebski A, Wyrzykowska P, Koj A, Jura J. Regulatory feedback loop between NF-κB and MCP-1-induced protein 1 RNase. FEBS J (2009) 276:5892–5905. doi:10.1111/j.1742-4658.2009.07273.x

48. Liang J, Saad Y, Lei T, Wang J, Qi D, Yang Q, Kolattukudy PE, Fu M. MCP-induced protein 1 deubiquitinates TRAF proteins and negatively regulates JNK and NF-κB signaling. J Exp Med (2010) 207:2959–2973. doi:10.1084/jem.20092641

49. Diamond MS, Farzan M. The broad-spectrum antiviral functions of IFIT and IFITM proteins. Nat Rev Immunol (2013) 13:46–57. doi:10.1038/nri3344

50. Buchanan EL, Mcalexander MA, Witwer KW. SAMHD1 expression in blood cells of HIV-1 elite suppressors and viraemic progressors. J Antimicrob Chemother (2015) 70:954–956. doi:https://doi.org/10.1093/jac/dku428

51. Uehata T, Iwasaki H, Vandenbon A, Matsushita K, Hernandez-Cuellar E, Kuniyoshi K, Satoh T, Mino T, Suzuki Y, Standley DM, et al. Malt1-induced cleavage of regnase-1 in CD4(+) helper T cells regulates immune activation. Cell (2013) 153:1036–49. doi:10.1016/j.cell.2013.04.034

52. Liang J, Song W, Tromp G, Kolattukudy PE, Fu M. Genome-wide survey and expression profiling of CCCH-zinc finger family reveals a functional module in macrophage activation. PLoS One (2008) 3: doi:10.1371/journal.pone.0002880

53. Matsushita K, Takeuchi O, Standley DM, Kumagai Y, Kawagoe T, Miyake T, Satoh T, Kato H, Tsujimura T, Nakamura H, et al. Zc3h12a is an RNase essential for controlling immune responses by regulating mRNA decay. Nature (2009) 458:1185–1190. doi:10.1038/nature07924

54. Liang J, Wang J, Azfer A, Song W, Tromp G, Kolattukudy PE, Fu M. A novel CCCH-zinc finger protein family regulates proinflammatory activation of macrophages. J Biol Chem (2008) 283:6337–46. doi:10.1074/jbc.M707861200

55. Li M, Cao W, Liu H, Zhang W, Liu X, Cai Z, Guo J, Wang X, Hui Z, Zhang H, et al. MCPIP1 down-regulates IL-2 expression through an ARE-independent pathway. PLoS One (2012) 7:e49841. doi:10.1371/journal.pone.0049841

56. Feinberg MW, Cao Z, Wara AK, Lebedeva MA, Senbanerjee S, Jain MK. Kruppel-like factor 4 is a mediator of proinflammatory signaling in macrophages. J Biol Chem (2005) 280:38247–58. doi:10.1074/jbc.M509378200

57. Feinberg MW, Wara AK, Cao Z, Lebedeva MA, Rosenbauer F, Iwasaki H, Hirai H, Katz JP, Haspel RL, Gray S, et al. The Kruppel-like factor KLF4 is a critical regulator of monocyte *A.*differentiation. EMBO J (2007) 26:4138–48. doi:10.1038/sj.emboj.7601824

58. Kapoor N, Niu J, Saad Y, Kumar S, Sirakova T, Becerra E, Li X, Kolattukudy PE. Transcription factors STAT6 and KLF4 implement macrophage polarization via the dual catalytic powers of MCPIP. J Immunol (2015) 194:6011–23. doi:10.4049/jimmunol.1402797

59. Yoon HS, Chen X, Yang VW. Kruppel-like factor 4 mediates p53-dependent G1/S cell cycle arrest in response to DNA damage. J Biol Chem (2003) 278:2101–5. doi:10.1074/jbc.M211027200

60. Zhang W, Geiman DE, Shields JM, Dang DT, Mahatan CS, Kaestner KH, Biggs JR, Kraft AS, Yang VW. The gut-enriched Kruppel-like factor (Kruppel-like factor 4) mediates the transactivating effect of p53 on the p21(WAF1)/(Cip)1 promoter. J Biol Chem (2000) 275:18391–18398. doi:10.1074/jbc.C000062200

61. Garg A, Kaul D, Chauhan N. APOBEC3G governs to ensure cellular oncogenic transformation. Blood Cells Mol Dis (2015) 55:248–54. doi:10.1016/j.bcmd.2015.07.009

62. Pauls E, Ruiz A, Riveira-Munoz E, Permanyer M, Badia R, Clotet B, Keppler OT, Ballana E, Este JA. p21 regulates the HIV-1 restriction factor SAMHD1. Proc Natl Acad Sci (2014) 111:E1322–E1324. doi:10.1073/pnas.1322059111

63. Allouch A, David A, Amie SM, Lahouassa H, Chartier L, Margottin-Goguet F, Barre-Sinoussi F, Kim B, Saez-Cirion A, Pancino G. p21-mediated RNR2 repression restricts HIV-1 replication in macrophages by inhibiting dNTP biosynthesis pathway. Proc Natl Acad Sci (2013) 110:E3997–E4006. doi:10.1073/pnas.1306719110

64. Leng J, Ho H-P, Buzon MJ, Pereyra F, Walker BD, Yu XG, Chang EJ, Lichterfeld M. A Cell-Intrinsic Inhibitor of HIV-1 Reverse Transcription in CD4+ T Cells from Elite Controllers. Cell Host Microbe (2014) 15:717–728. doi:10.1016/j.chom.2014.05.011

65. Wang D, de la Fuente C, Deng L, Wang L, Zilberman I, Eadie C, Healey M, Stein D, Denny T, Harrison LE, et al. Inhibition of Human Immunodeficiency Virus Type 1 Transcription by Chemical Cyclin-Dependent Kinase Inhibitors. J Virol (2001) 75:7266–7279. doi:10.1128/JVI.75.16.7266-7279.2001

66. Kumari N, Iordanskiy S, Kovalskyy D, Breuer D, Niu X, Lin X, Xu M, Gavrilenko K, Kashanchi F, Dhawan S, et al. Phenyl-1-Pyridin-2yl-Ethanone-Based Iron Chelators Increase IκB-α Expression, Modulate CDK2 and CDK9 Activities, and Inhibit HIV-1 Transcription. Antimicrob Agents Chemother (2014) 58:6558–6571. doi:10.1128/AAC.02918-14

67. Lin R-J, Chien H-L, Lin S-Y, Chang B-L, Yu H-P, Tang W-C, Lin Y-L. MCPIP1 ribonuclease exhibits broad-spectrum antiviral effects through viral RNA binding and degradation. Nucleic Acids Res (2013) 41:3314–3326. doi:10.1093/nar/gkt019

68. Arias CF, Ballesteros-Tato A, Garcia MI, Martin-Caballero J, Flores JM, Martinez-A C, Balomenos D. p21CIP1/WAF1 Controls Proliferation of Activated/Memory T Cells and Affects Homeostasis and Memory T Cell Responses. J Immunol (2007) 178:2296–2306. doi:10.4049/jimmunol.178.4.2296

69. Li Y, Huang X, Huang S, He H, Lei T, Saaoud F, Yu X-Q, Melnick A, Kumar A, Papasian CJ, et al. Central role of myeloid MCPIP1 in protecting against LPS-induced inflammation and lung injury. Signal Transduct Target Ther (2017) 2:17066. doi:10.1038/sigtrans.2017.66

70. Uehata T, Takeuchi O. Regnase-1 Is an Endoribonuclease Essential for the Maintenance of Immune Homeostasis. J Interf Cytokine Res (2017) 37:220–229. doi:10.1089/jir.2017.0001

71. De Pablo A, Bogoi R, Bejarano I, Toro C, Valencia E, Moreno V, Martín-Carbonero L, Gómez-Hernando C, Rodés B. Short communication: p21/CDKN1A expression shows broad interindividual diversity in a subset of HIV-1 elite controllers. AIDS Res Hum Retroviruses (2016) 32:1–5. doi:10.1089/aid.2015.0137

72. Abdel-Mohsen M, Wang C, Strain MC, Lada SM, Deng X, Cockerham LR, Pilcher CD, Hecht FM, Liegler T, Richman DD, et al. Select host restriction factors are associated with HIV persistence during antiretroviral therapy. AIDS (2015) 29:411–20. doi:10.1097/QAD.0000000000000572

73. Lin R-J, Chu J-S, Chien H-L, Tseng C-H, Ko P-C, Mei Y-Y, Tang W-C, Kao Y-T, Cheng H-Y, Liang Y-C, et al. MCPIP1 suppresses hepatitis C virus replication and negatively regulates virus-induced proinflammatory cytokine responses. J Immunol (2014) 193:4159–68. doi:10.4049/jimmunol.1400337

