## Supplementary Material for "Increased expression of MCPIP1 in HIV-1 controllers is correlated with overexpression of p21"

***Correspondence:**

Suwellen S. D. de Azevedo

### Supplementary Figures

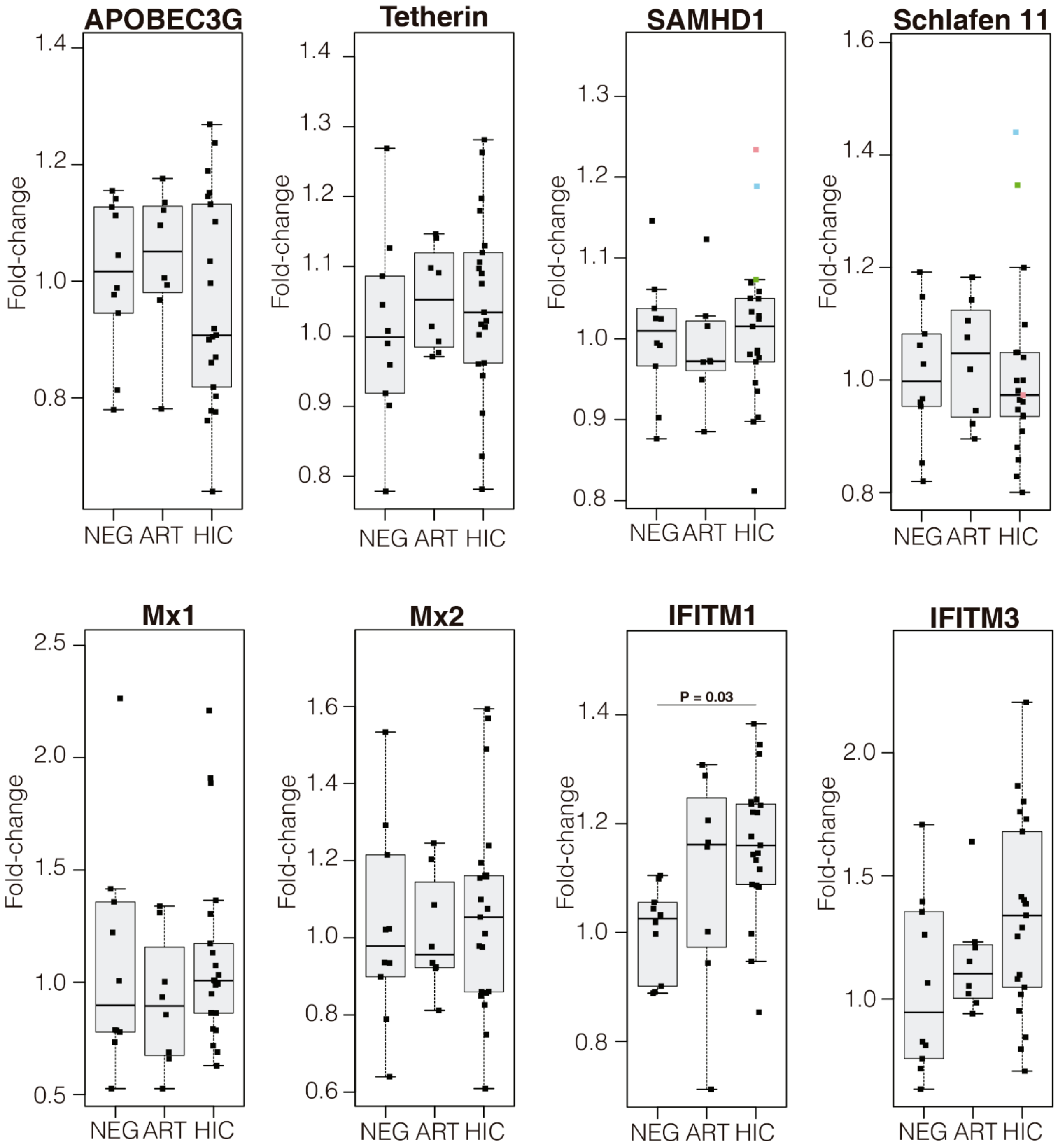


**Supplementary Figure S1**. Boxplots represent the interquartile and sample median (central solid black line) of the relative changes (fold-change values relative to the mean of HIV-1-uninfected (NEG) subjects) of different restriction factor comparing NEG and ART-suppressed subjects (ART) with HIV controllers (HIC). The RF's names used in the analysis are indicated above each graph. HIC exhibited statistically significant differences (P-values < 0.05) with respect to NEG group only for IFITM1. The colored squares (one per individual) in SAMHD1 and Schlafen 11 from HIC represent individuals with mRNA levels well above the normal range in one or both RF.

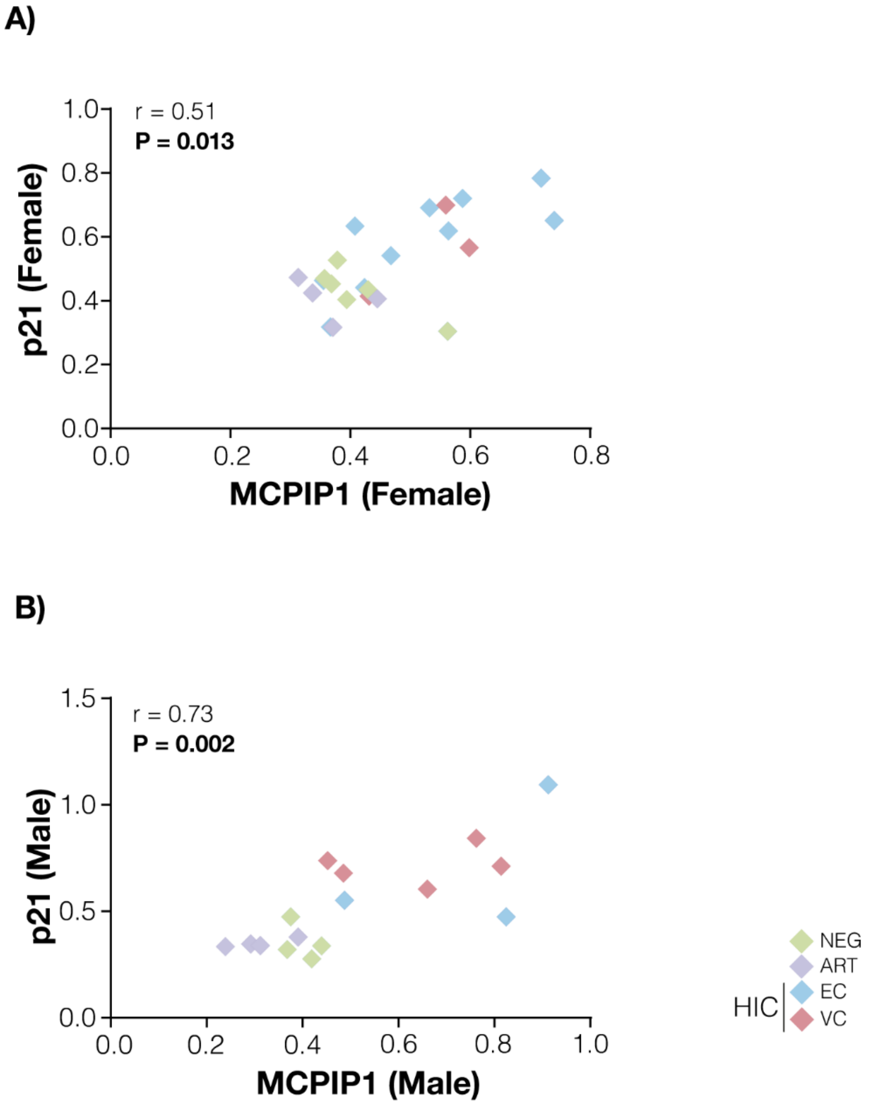


**Supplementary Figure S2**. MCPIP1 and p21 mRNA levels in PBMC are positively correlated, regardless of sex. The MCPIP1 and p21 normalized expression correlations were calculated considering all groups. The points' colors indicate the patient group, accordingly to the legend. Correlation coefficients (Spearman's ρ) are shown in the upper left corner of each graph. P-values $< 0.05$ were considered statistically significant.

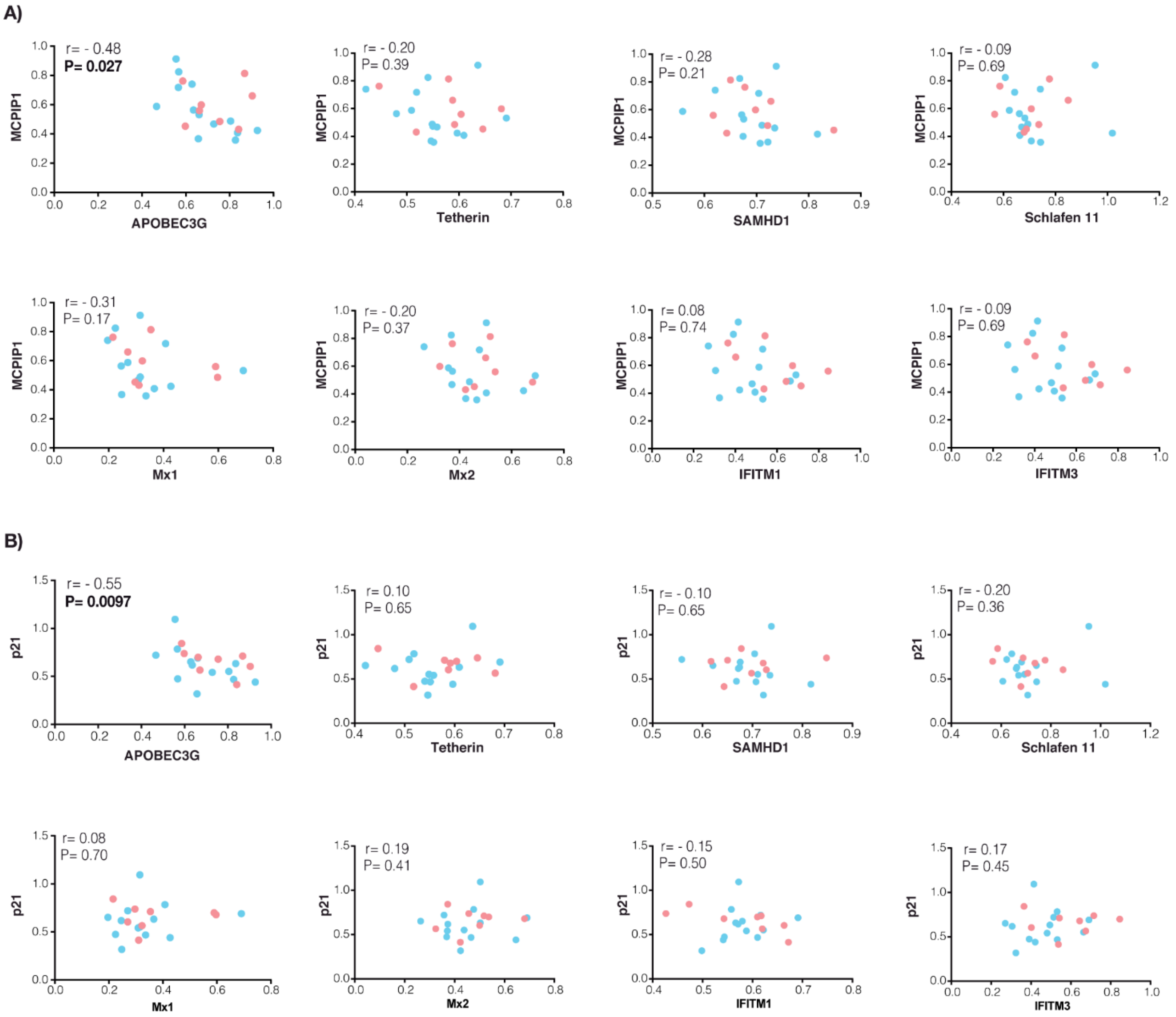


**Supplementary Figure S3.** Correlations between normalized expression levels of MCPIP1 (A) and p21 (B) with several anti-HIV-1 restriction factors (RF) in HIC. Blue points represent values form elite controllers while the red ones represent values from viremic controllers. The RF's names used in the correlation are indicated on the x-axis and the corresponding correlation coefficient (Spearman's rho) are shown in each graph.

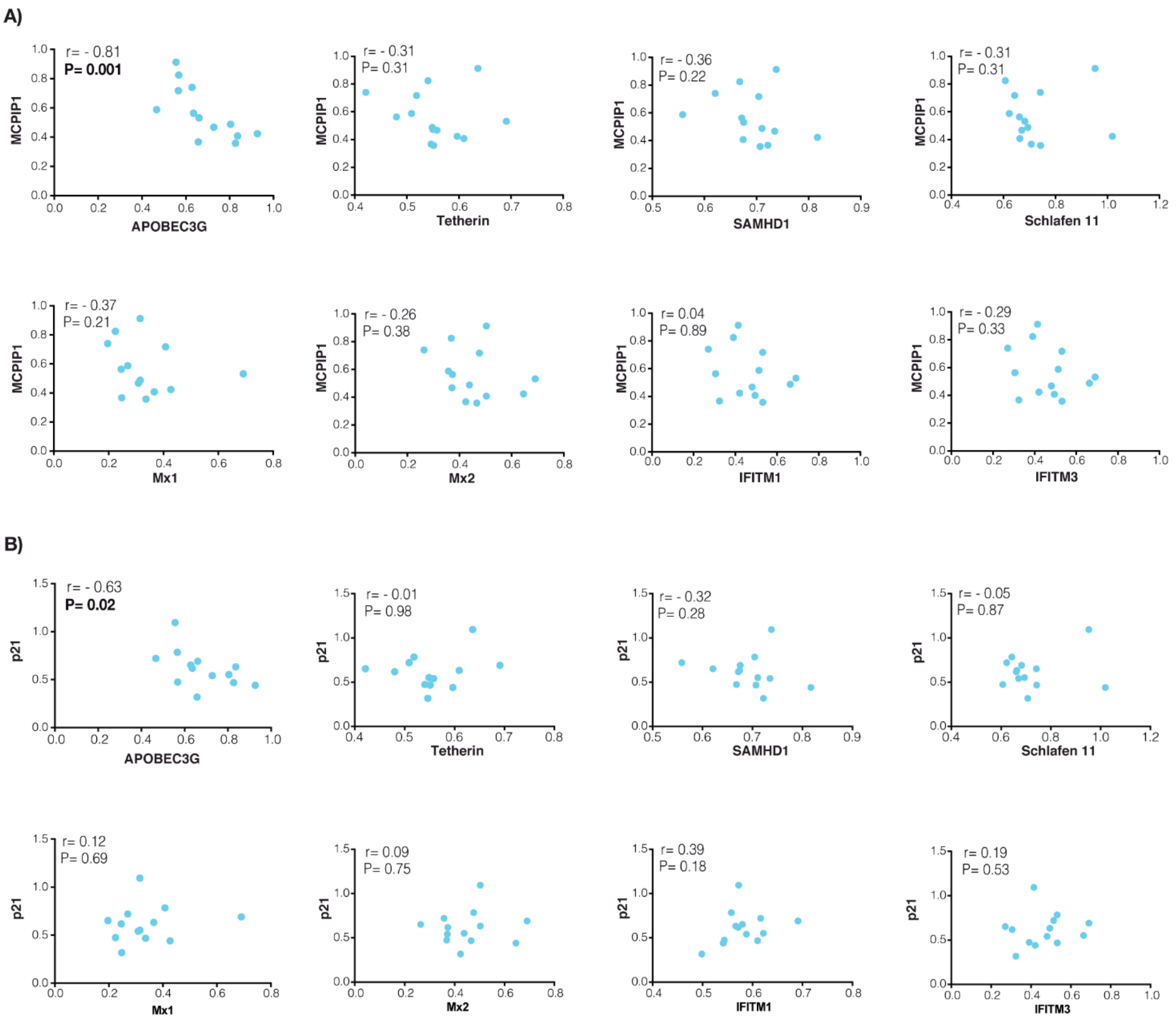


**Supplementary Figure S4.** Correlations between normalized expression levels of MCPIP1 (A) and p21 (B) with several anti-HIV-1 restriction factors (RF) in EC. Blue points represent values form elite controllers while the red ones represent values from viremic controllers. The RF's names used in the correlation are indicated on the x-axis and the corresponding correlation coefficient (Spearman's rho) are shown in each graph.

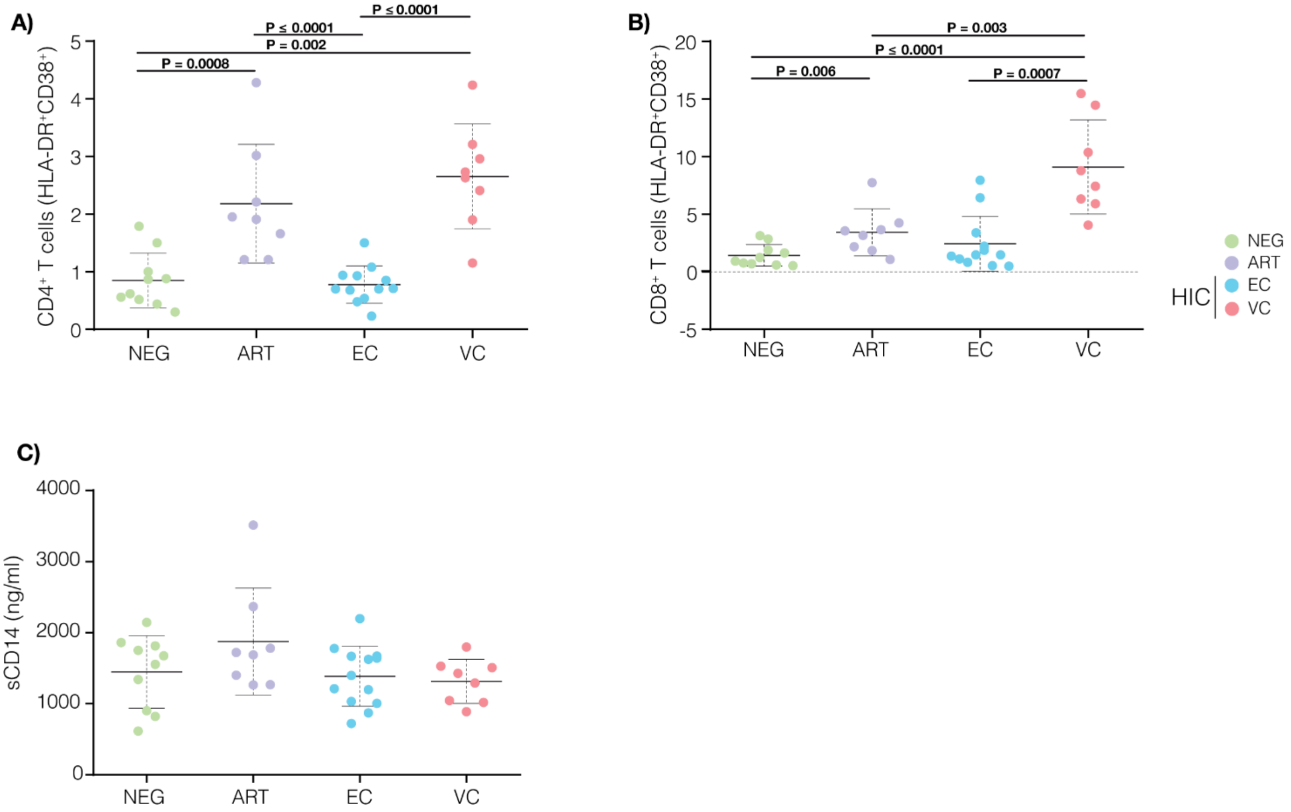


**Supplementary Figure S5.** The mean of activated CD4$^+$ T cells (HLA-DR$^+$CD38$^+$) counts (A), activated CD8$^+$ T cells (HLA-DR$^+$CD38$^+$) counts (B), and soluble CD14 (sCD14) in plasma (C) were compared for each group. The color of each dot represents the group as indicated in the legend at right. P-values < 0.05 were considered statistically significant.

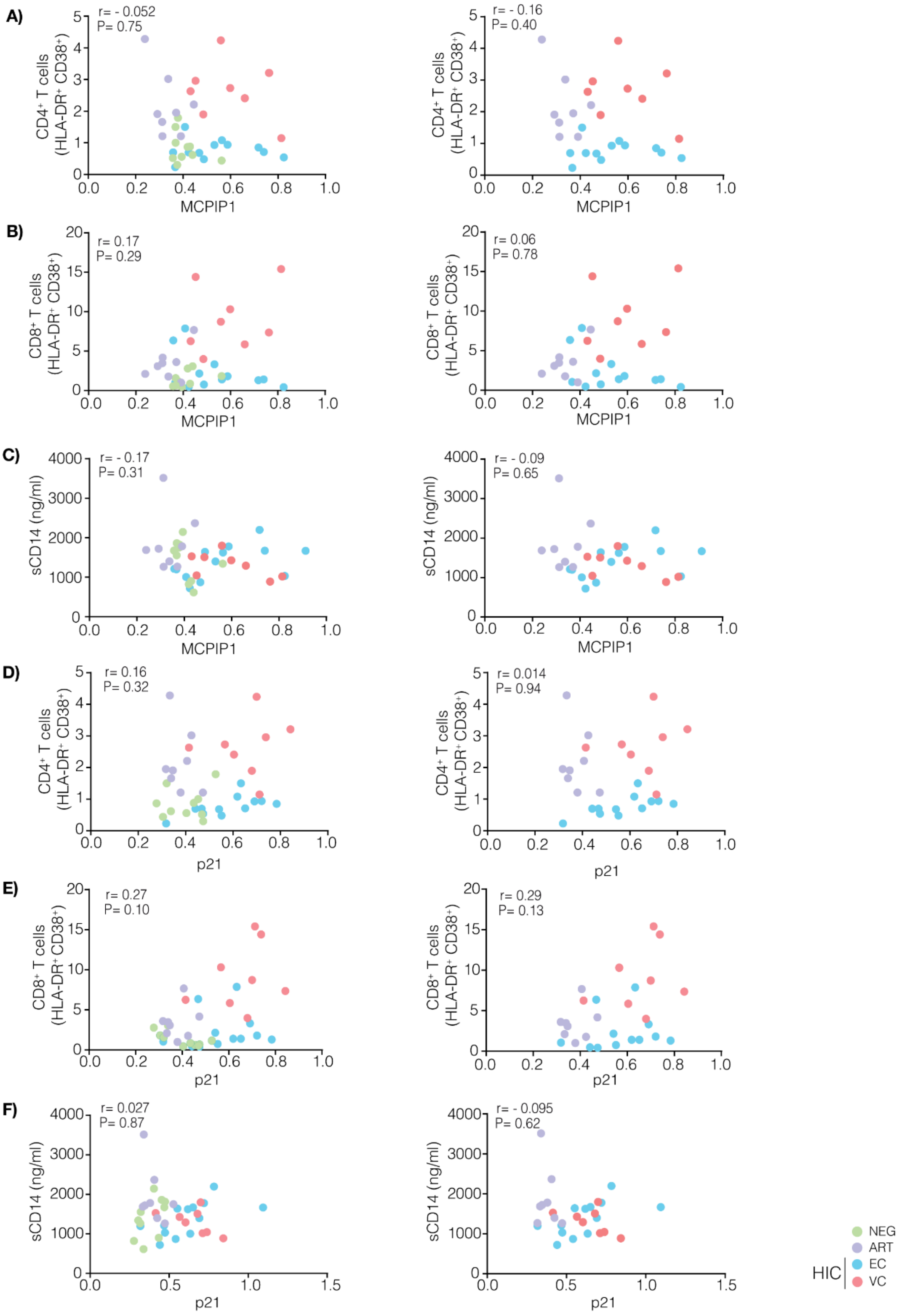


**Supplementary Figure S6**. MCPIP1 and p21 are not correlated with CD4$^+$/CD8$^+$ T cell, and monocyte activation in all groups and in HIV-1 infected individuals. Correlations were made evaluating the relationship between activated CD4$^+$, CD8$^+$ T cells or sCD14 levels with the normalized expression of MCPIP1 (A, B, and C, respectively) and p21 (D, E and F, respectively) for different combinations of groups. The points' colors present in each graph indicate the groups present according to the legend. The correlation coefficient (Spearman's rho) are shown in the left corner.

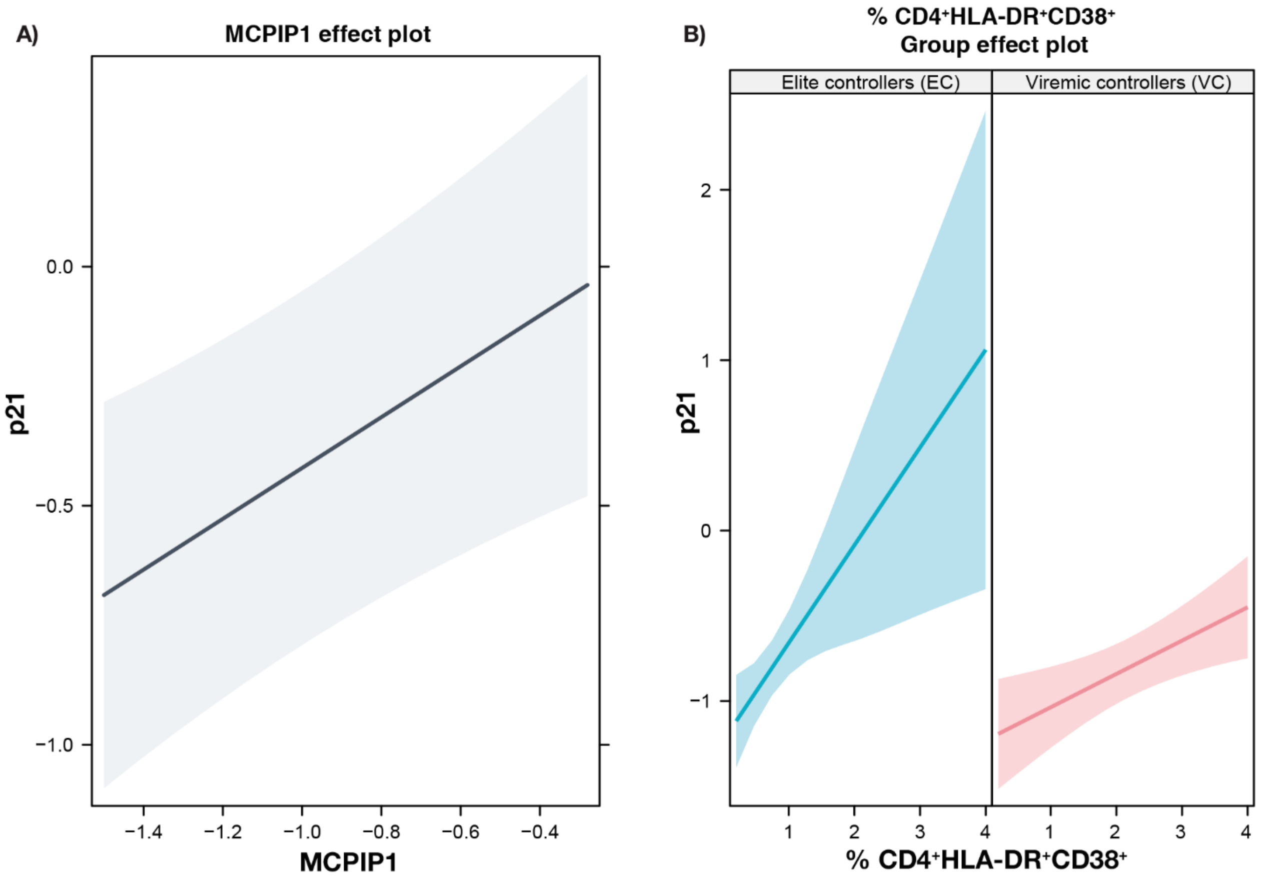


**Supplementary Figure S7**. MCPIP1 and the frequency of $CD4^{+}HLA\text{-}DR^{+}CD38^{+}$ T cells up-regulate p21 mRNA levels in PBMC from HIC. Effects plots demonstrating (A) the up-regulation of MCPIP1 is positively associated with the increase of the expression of p21 in PBMC from HIC; while in (B) the frequency of $CD4^{+}HLA\text{-}DR^{+}CD38^{+}$ T cells is positively associated with the increase of the expression of p21 in PBMC from both elite (EC) and viremic (VC) controllers, and this increase was of the p21 expression was down-regulated by the increase of $CD4^{+}HLA\text{-}DR^{+}CD38^{+}$ T cells in VC when compared to EC individuals. P-values $< 0.05$ were considered statistically significant.

**Supplementary Table 1**. Distribution of individuals in different groups according to sex.

| Group | n (%) | Sex (n [%]) | | *P- value** |
|---|---|---|---|---|
| | | Female | Male | |
| NEG | 10 (25.6) | 6 (15.4) | 4 (10.3) | 0.9083 |
| ART | 8 (20.5) | 4 (10.3) | 4 (10.3) | |
| HIC | 21 (53.8) | 13 (33.3) | 8 (20.5) | |
| NEG | 10 (25.6) | 6 (15.4) | 4 (10.3) | 0.3273 |
| ART | 8 (20.5) | 4 (10.3) | 4 (10.3) | |
| EC | 13 (33.3) | 10 (25.6) | 3 (7.7) | |
| VC | 8 (20.5) | 3 (7.7) | 5 (12.8) | |

* Statistical analyses were performed using the Fisher exact test.
